# Circadian regulation of light-evoked attraction/avoidance in day- vs. night-biting mosquitoes

**DOI:** 10.1101/627588

**Authors:** Lisa Soyeon Baik, Ceazar Nave, David D. Au, Tom Guda, Joshua A. Chevez, Anandasankar Ray, Todd C. Holmes

## Abstract

Mosquitoes pose widespread threats to humans and other animals as disease vectors. Day- vs. night-biting mosquitoes occupy distinct time-of-day niches and exhibit very different innate temporal attraction/avoidance behavioral responses to light, yet little is known about their circuit or molecular mechanisms. Day-biting diurnal mosquitoes *Aedes aegypti* are attracted to light during the day regardless of spectra. In contrast, night-biting nocturnal mosquitoes *Anopheles coluzzii* avoid short, but not long wavelength light. Attraction/avoidance behavioral responses to light in both species change with time-of-day and show distinct sex and circuit differences. The basis of diurnal versus nocturnal behavior is driven by clock timing, which cycle anti-phase between day-biting versus night-biting mosquito species. Disruption of the circadian molecular clock severely interferes with light-evoked attraction/avoidance behavior in mosquitoes. In summary, attraction/avoidance mosquito behaviors are circadian and light regulated, which may be applied towards species specific control of harmful mosquitoes.

## Introduction

Mosquitoes present a worldwide threat as key disease vectors that spread malaria parasites and Zika, Chikungunya, West Nile, Yellow Fever, and Dengue Fever viruses. Toxic pesticides are environmentally costly in contrast to light-based insect control approaches. However, light-based insect control approaches do not typically take into consideration the day- vs. night behavioral activity profiles of insects. Insects display a wide range of short wavelength light modulated behaviors, including attraction/avoidance^1-7^. It has been long assumed that insect responses to ultraviolet (UV) light are mediated by UV-sensitive opsins expressed in eyes and other external photoreceptors. Mosquitoes and flies also express non-opsin photoreceptors including the blue-light sensitive flavoprotein CRYPTOCHROME (CRY)^8^ which mediates a wide range of behavioral responses to blue and UV light, including circadian modulated attraction and avoidance^1,2^. Different mosquito species have evolved distinct circadian timing of behaviors according to their temporal/ecological niches, which likely minimize inter-species competition. Some mosquito species are diurnal (i.e., *Aedes aegypti*) while others are nocturnal (i.e., *Anopheles coluzzii*). Numerous mosquito behaviors change with the time-of-day, including flight activity, mating, oviposition, and biting^9-15^. Considering their impact on health and ecology, little is known about the species basis of diurnality/nocturnality and behavioral timing in mosquitoes.

## Results

We measured the light environment preference behaviors of diurnal (*Aedes aegypti*) and nocturnal (*Anopheles coluzzii*) mosquito species throughout the 24 hr day using a custom designed arena (S1 Fig.). Young adult mosquitoes were presented with a choice of light versus shaded environments during the subjective daytime (ZT 0-12) to measure preference for the light-exposed or in the shaded-environment, quantified as % of preference. Nocturnal versus diurnal mosquito species exhibit striking differences in their light-evoked attraction/avoidance behavior, despite constant light intensity. Diurnal *Aedes aegypti* (*Ae. aegypti*) females are behaviorally attracted to UV light during the day (Fig. 1A and 1J). In contrast, nocturnal *Anopheles coluzzii* (*An. coluzzii)* females strongly avoid UV light during most of the daytime (Fig. 1B and 1J). Both *Ae. aegypti and An. coluzzii* females show shifted attraction/avoidance behavior as dusk approaches even under constant light intensity until lights off (Fig. 1A, 1B, 1E, and 1G). This temporal “anticipatory” afternoon behavioral shift starts much earlier for diurnal *Ae. aegypti* females, which increase their attraction behavior beginning around mid-afternoon (ZT6) (Fig. 1A and 1E). In contrast, nocturnal *An. coluzzii*, females show sharp decreases in UV light avoidance starting before dusk (ZT11) (Fig. 1G). Precisely at dusk, they enter the previously UV illuminated area, rapidly reaching 60% preference for this area 10 minutes after dusk and peak 70% preference one hour after dusk (Fig. 1B).

**Figure 1.**
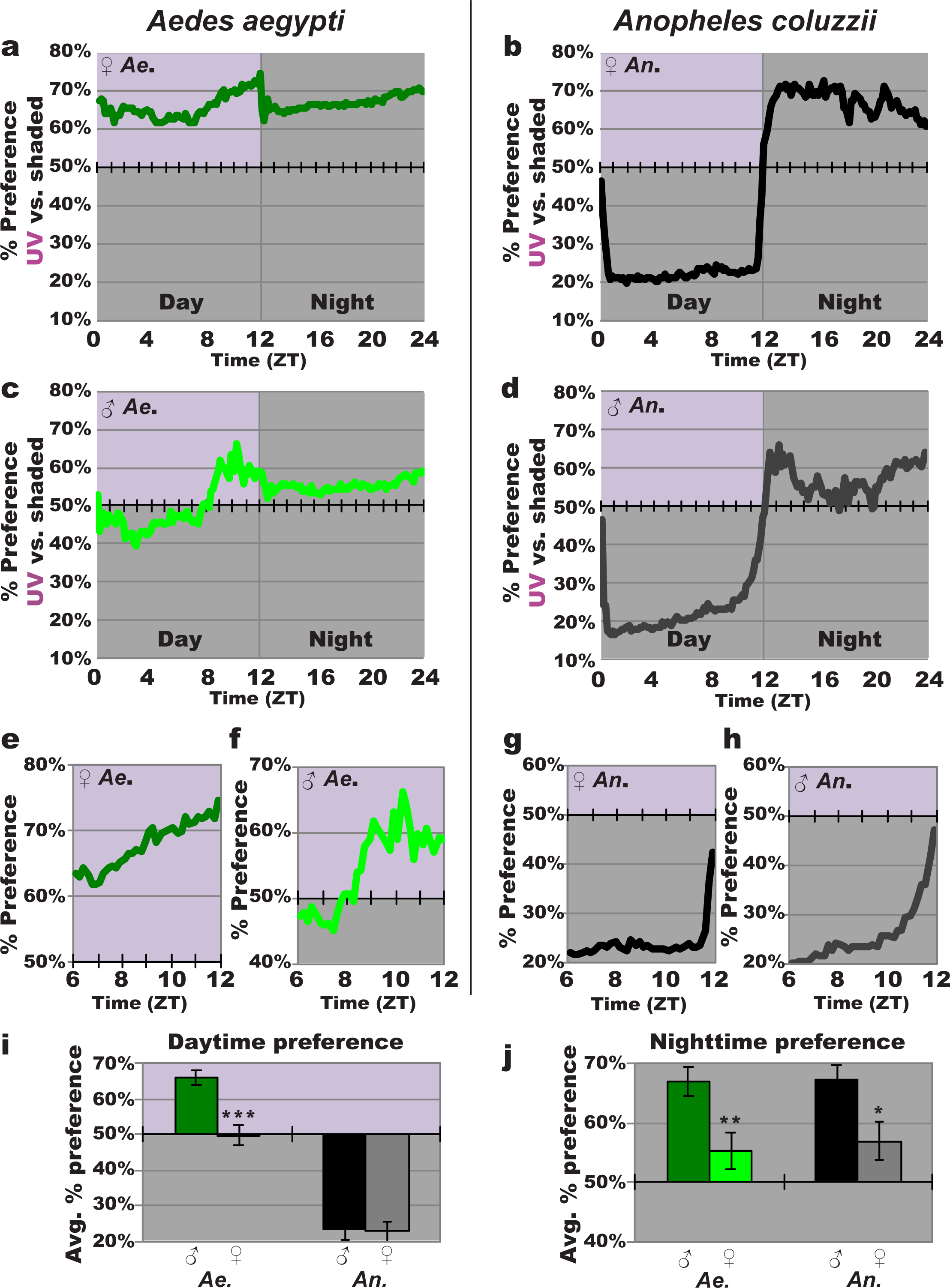
UV light-evoked attraction/avoidance responses in diurnal and nocturnal mosquitoes are specie- and sex-dependent. **(a-d)** Attraction/avoidance behavior to UV light, measured by % of preference in UV-exposed versus shaded environment throughout 12 hr: 12 hr UV light: dark for female *Ae. aegypti* (n=110) **(a)**, female *An. coluzzii* (n=88) **(b)**, male *Ae. aegypti* (n=61) **(c)**, male *An. coluzzii* (n=47) **(d). (e-h)** Attraction/avoidance behavior to UV light, measured by % of preference in UV-exposed versus shaded-environment during ZT 6-12 for female *Ae. aegypti* **(e)**, male *Ae. aegypti* **(f)**, female *An. coluzzii* **(g)**, male *An. coluzzii* **(h). (i-j)** Average attraction/avoidance behavioral preference to UV light-exposed versus shaded-environment for daytime **(i)** and nighttime **(j)** in *Ae. aegypti* and *An. coluzzii* female and male mosquitoes. Data are represented as mean ± S.E.M. *p < 0.05; **p < 0.01; ***p < 0.001 vs. female.

Male mosquitoes form swarms in anticipation of females, thus we considered the possibility of sex differences for avoidance/attraction behavioral responses to UV light for both diurnal and nocturnal mosquito species. Diurnal *Ae. aegypti* males are attracted to UV light during the late subjective daytime, but to a significantly lesser extent than females, which are attracted to UV light throughout the entire day (Fig. 1C and 1I). Nocturnal *An. coluzzii* males strongly avoid UV light, similar to *An. coluzzii* females (Fig. 1D and 1I). Both species show sex-specific differences in timing of “anticipation” of increase in attraction/decrease in avoidance while temporally approaching dusk. *Ae. aegypti* male attraction peaks much earlier (ZT10) than that of females (ZT 12) (Fig. 1A, 1C, 1E, and 1F). Similarly as dusk approaches, *An. coluzzii* males initiate decreases in avoidance much earlier than females during their transition to attraction to the previously UV light illuminated area (Fig. 1B, 1D, 1G, and 1H). Sex-dependent differences persist even after the UV light is turned off, simulating the subjective nighttime (ZT 12-24). For both species, females prefer the previously light-exposed environment throughout the night. This nighttime preference for previously UV-light exposed environment is significantly higher in females, compared to males (Fig. 1A-1D, and 1J).

The color spectral specificity of attraction/avoidance behavior varies between different insect species^1,2,4,5,16^. *Drosophila melanogaster* diptera fruit flies avoid short wavelength light, but not long wavelength light and *Drosophila* UV light avoidance peaks in the midday during their low locomotor activity “siesta”^1,2^. To determine the spectral dependence of mosquito attraction/avoidance behavioral light responses, we compared their environmental preferences for visible short wavelength blue light and visible long wavelength red light for comparison with UV light (Fig. 1). Diurnal *Ae. aegypti* females are attracted to both blue and red light during the daytime, very similar to their behavioral attraction to UV light (Fig. 1A, and Panels A, C, and E in S2 Fig.). *Ae. aegypti* females are equally attracted to all light wavelengths tested (Panel E in S2 Fig.). Nocturnal *An. coluzzii* females, which strongly avoid UV light (Fig. 1B), also avoid blue light during the day (Panel B in S2 Fig.). Their magnitude of blue light avoidance is significantly lower than their UV light avoidance (Panel E in S2 Fig.). In contrast to their striking short wavelength avoidance during the day, female *An. coluzzii* are slightly attracted to long wavelength red light (Panels D and E in S2 Fig.). During the nighttime, females of both species prefer environments with prior UV light-exposure (Fig. 1J and Panel F in S2 Fig.), significantly higher than their nighttime preference for areas with prior blue- or red-light exposure, for which they are weakly attracted to (Panel F in S2 Fig.). Thus, attraction/avoidance behavioral responses are wavelength-dependent and differ in both overall profile and anticipation of dusk between nocturnal and diurnal species.

We anatomically mapped the circadian neuronal network in the central brain of female diurnal and nocturnal mosquitoes, motivated by the apparent relationship between the circadian clock and circuit modulation of light attraction/avoidance behavior, defined by the cyclic expression of PERIOD (PER) clock protein that drives rhythmic changes in physiology and behavior. In *Drosophila melanogaster*, Pigment Dispersing Factor (PDF) is a neuropeptide co-expressed with PER in the small- and large-lateral ventral neurons (LNvs), which modulate circadian- and light-mediated behaviors such as circadian locomotion, sleep, arousal, and light-evoked attraction/avoidance behaviors^1,2,17-20^. PER and PDF proteins are co-expressed in a similar neuroanatomical pattern the lateral ventral area in both *Ae. aegypti* and *An. coluzzii* female adult brains and are anatomically similar to *Drosophila* melanogaster and some other insects^21,22^. These PDF^+^ and PER^+^ neurons can be further distinguished by size, location and projections as large- (l-LNvs) and small-lateral ventral neurons (s-LNvs) in *Ae. aegypti* and *An. coluzzii* (Fig. 2 and S3 Fig.) and feature large neuronal arbors in the optic lobes dorsal projections, similar to *Drosophila* ^20^ (S4 Fig., S1 Movie, and S2 Movie). The neuroanatomical locations of PER^+^ neurons show species specific similarities and differences for cell groups between diurnal *Ae. aegypti* and nocturnal *An. coluzzii* outside of the lateral ventral area. Similarly located neuronal groups include putative dorsal neurons (DNs) in *Ae. aegypti* and *An. coluzzii* (Fig. 2 and S3 Fig.). Differentially located neuronal groups include approximately 5 PER^+^ neurons in the medial-anterior region of *Ae. aegypti* female brains, which we call medial-anterior neurons (m-ANs) here (Fig. 2 and S3 Fig.). Based on location and size, another differentially located neuronal group include approximately 7 PER^+^/PDF^−^ neurons, which resemble the pars intercerebralis (PI) in *An. Coluzzii* (Fig. 2). In *Drosophila melanogaster*, PI neurons are physiologically circadian rhythmic although by way of other clock neuron input^23,24^. These PI-like PER^+^/PDF^−^ neurons are not detected in *Ae. aegypt* (Fig. 2 and S3 Fig.). In summary, we observe both shared and distinct anatomical features of circadian neuronal circuit of diurnal versus nocturnal mosquito species.

**Figure 2.**
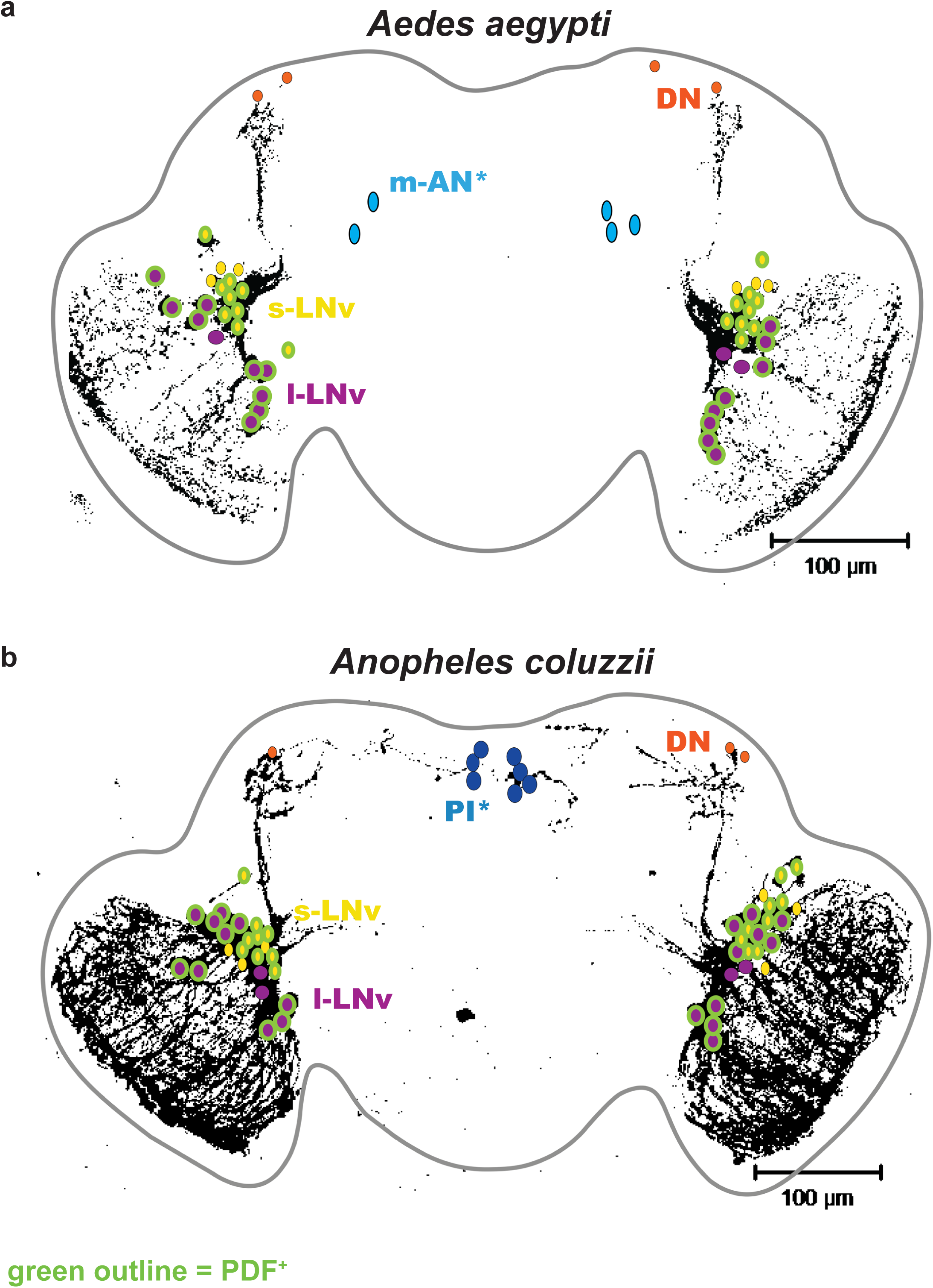
Schematic representation of *Aedes aegypti* and *Anopheles coluzzii* circadian neuronal circuits. **(a-b)** Illustration of representative adult female central brains and its neurons expression PER and/or PDF. Asterisk (*) indicates groups distinct for each species. **(a)** *Ae. aegypti* s-LNv in color yellow, l-LNv in color violet, DNs in color orange, and m-AN in color light blue, with PDF^+^ neurons indicated with green outline. **(b)** *An. coluzzii* s-LNv in color yellow, l-LNv in color violet, DNs in color orange, and PI neurons in color dark blue, with PDF^+^ neurons indicated with green outline.

We measured PER protein oscillating levels throughout the 24hr day using anti-PER immunocytochemistry staining, co-stained with PDF, at 6hr intervals in the brains of standard 12hr: 12hr light:dark (LD) entrained *Ae. aegypti* and *An. coluzzii* female mosquitoes. PER rhythms cycle robustly in both *Ae. aegypti* and *An. coluzzii* circadian neurons in a neuronal subgroup-specific manner. Notably, PER cycling in PDF^+^ LNvs oscillate in opposite phases between diurnal *Ae. aegypti* versus nocturnal *An. coluzzii* brains (Fig. 3). PER protein levels peak in late night/early day in PDF^+^ s-LNv and l-LNv of the diurnal mosquito *Ae. aegypti* (Fig. 3A, 3C, and 3D). In contrast, PER protein levels peak in late day/early night in PDF^+^ s-LNv and l-LNv of the nocturnal mosquito *An. coluzzii* (Fig. 3B, 3E, and 3F). PER protein levels peak in early daytime in PER^+^ DNs of both diurnal *Ae. aegypti* and nocturnal *An. coluzzii* (S5 Fig. and S7 Fig.). Similarly, PER protein levels peak during the daytime in *Ae. aegypti* specific m-ANs (S6 Fig.). In *An. coluzzii* PI-like neurons, PER protein levels peak in the early day (S8 Fig.). In summary, diurnal and nocturnal mosquitoes have distinct circadian molecular signatures in the brain, with an early day PER peak in diurnal mosquito *Ae. aegypti* versus an early evening PER peak in nocturnal *An. coluzzii* in PDF^+^ LNv neurons. These opposing oscillation phases suggest a mechanism for diurnal and nocturnality.

**Figure 3.**
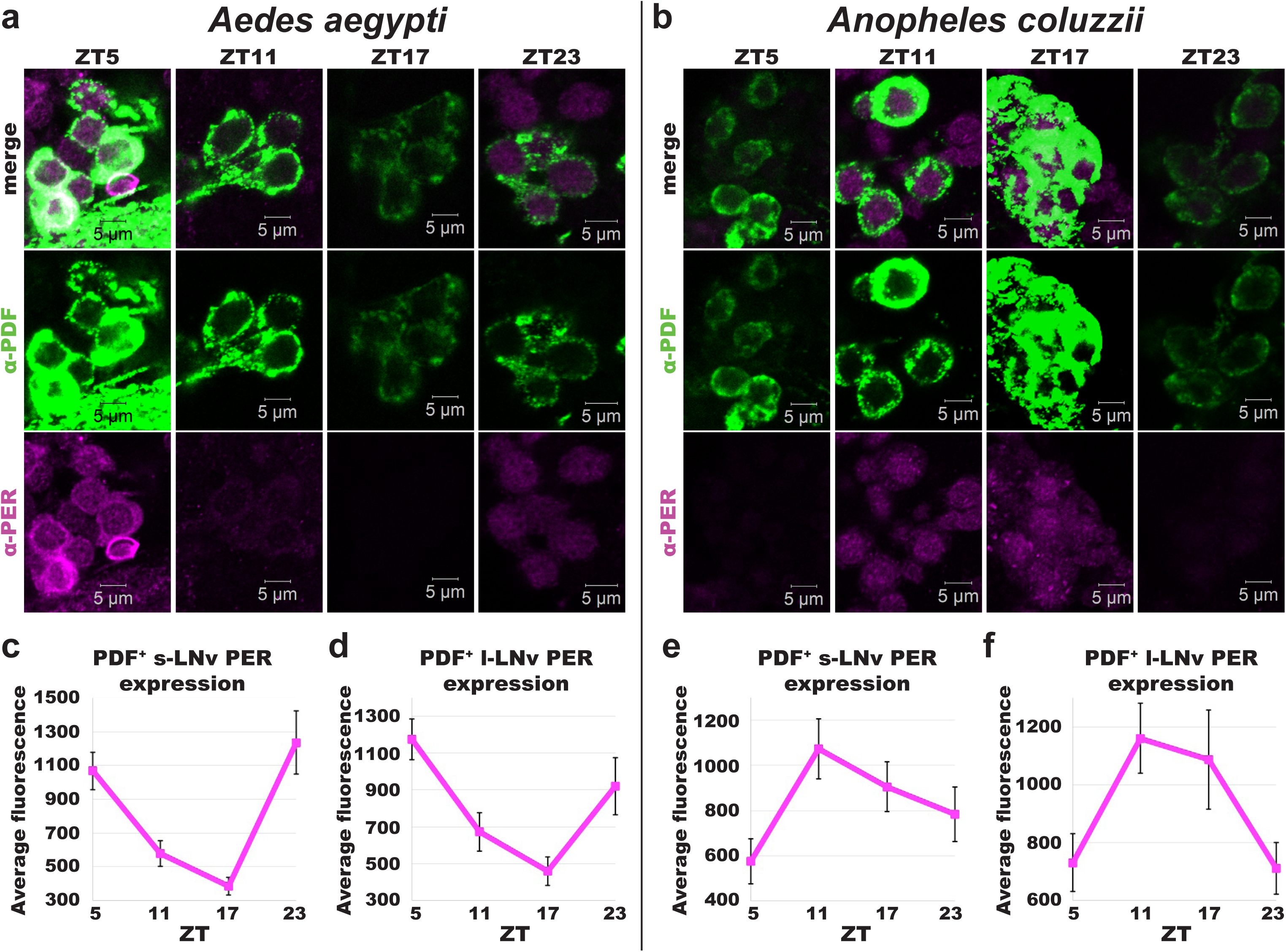
Diurnal versus nocturnal mosquito PER expression in PDF^+^ LNv neurons oscillate in anti-phasic manner. **(a-b)** Representative confocal images of adult female *Ae. aegypti* **(a)** and *An. coluzzii* **(b)** mosquito lateral ventral neurons (LNv) immunocytochemistry stained with α–PER (magenta) and α–PDF (green) antibodies at ZTs 5, 11, 17, and 23. **(c-f)** PERIOD expression levels at each ZT for *Ae. aegypti* (ZT5, n=27; ZT11, n=17; ZT17, n=6, ZT23, n=7) s-LNv **(c)** and l-LNv **(d)**, and *An. colluzzii* (ZT5, n=13; ZT11, n=31; ZT17, n=9, ZT23, n=8) s-LNv **(e)** and l-LNv **(f)**. Data are represented as mean ± S.E.M.

Constant light condition disrupts circadian clock gene expression and rhythmic behaviors in many animals, including mosquitoes^10,11,25^. To examine the functional linkage between circadian clock disruption and attraction/avoidance behavioral responses to UV light, *Ae. aegypti* and *An. coluzzii* female mosquitoes were exposed to constant UV light exposure (UV LL) for 3-5 days. Using anti-PER immunocytochemistry, we measured PER protein levels corresponding to species specific peak times in female mosquito brains following 3-5 days of UV LL. PER protein levels are severely reduced in mosquito brains following UV LL compared to LD in both *Ae. aegypti* and *An. coluzzii* (Fig. 4A-4D). In many brains, PER protein levels in LNvs could not be quantified because there was no visible PER staining following UV LL (not shown). We then measured the behavioral preference for UV-exposed versus shaded light environments under UV LL condition. During UV LL condition, both *Ae. aegypti* and *An. coluzzii* mosquitoes lack clear time-of-day dependent changes in attraction/avoidance behavior, including the anticipation (Fig. 4E-4H). *Ae. aegypti* females are attracted to UV light regardless of time-of-day (Fig. 4E and 4G). *An. coluzzii* females lack subjective day versus night differences in avoidance/attraction, and overall lack any clear preferences for either UV-exposed or shaded environments under UV LL condition (Fig. 4F and 4H). Clock ablation severely disrupts the timing of UV-evoked attraction/avoidance behavior of both diurnal and nocturnal mosquitoes.

**Figure 4.**
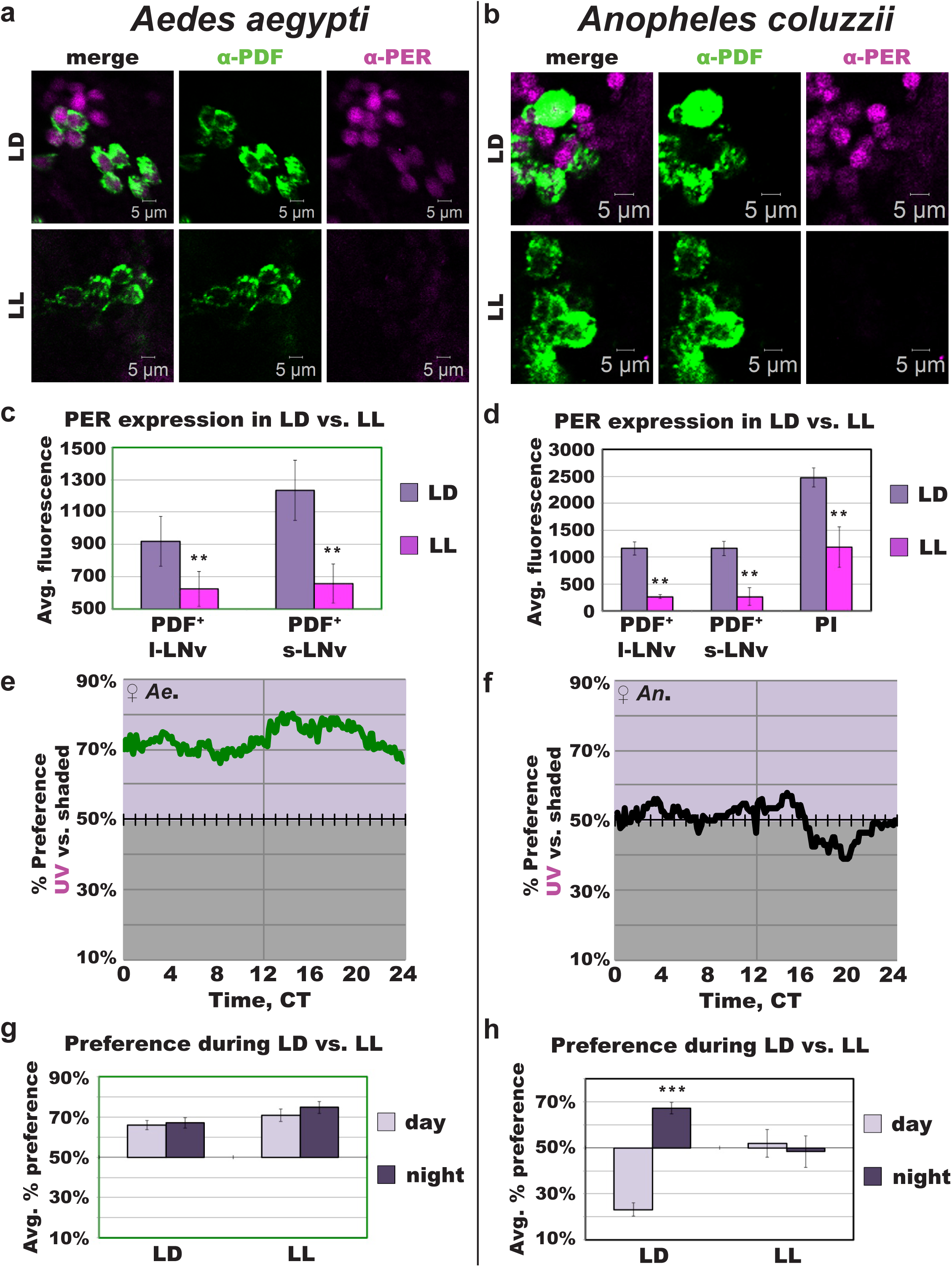
Constant UV light exposure disrupts circadian protein expression and clock modulation of attraction/avoidance behavioral responses to UV light in mosquitoes. **(a-b)** Representative confocal images of anti–PER (magenta) and anti–PDF (green) immunocytochemistry stained adult female mosquito brains under LD or following UV LL exposure for *Ae. aegypti* **(a)** and *An. coluzzii* **(b). (c-d)** Average fluorescence intensity of circadian neurons under LD versus LL conditions for *Ae. aegypti* ZT/CT 23 (LD, n=7; LL, n=13) **(c)** and *An. coluzzii* ZT/CT 11 (LD, n=31; LL, n=6) **(d). (e-f)** Attraction/avoidance behavior to UV light, measured by % of preference in UV-exposed versus shaded during UV LL for female *Ae. aegypti* **(e)**, female *An. coluzzii* **(f). (g-h)** Average attraction/avoidance behavioral preference to light-exposed versus shaded-environment during subjective daytime versus nighttime under LD or LL conditions for *Ae. aegypti* **(g)** and *An. coluzzii* **(h)** female mosquitoes. Data are represented as mean ± S.E.M. *p < 0.05; **p < 0.01; ***p < 0.001 vs. LD or day.

## Discussion

We show that mosquito attraction/avoidance behavior to light is specie-, sex-, spectra-, and time-of-day-dependent. Diurnal versus nocturnal mosquitoes have opposite attraction/avoidance behavioral valence for short wavelength light. Daytime-active mosquitoes, *Ae. aegypti*, are attracted to a wide range of light spectra during the daytime. In contrast nighttime-active mosquitoes *An. coluzzii* are strongly photophobic in response to short wavelength light. Both species exhibit anticipatory behavior of decreased avoidance and increased attraction during the temporal approach to dusk. This correlates with the ecological timing of increased flight activity and host seeking behaviors^10,11^. Interestingly, males of both diurnal and nocturnal species show earlier anticipatory behavior of increased UV attraction, compared to females. Both *Ae. aegypti* and *An. coluzzii* males exhibit earlier flight activity onset towards dusk, compared to females^15,26^. Male mosquitoes form ‘swarms’ in anticipation of female mosquitoes flying through to mate. The timing of male swarming is species specific. This dictates a temporal niche for mating of different mosquito species, whereby male mosquitoes evolved pre-dusk earlier anticipation to optimize their chance of mating^12,15^. Our previous work shows that light-evoked attraction/avoidance behavior is mediated by both opsin- and non-opsin based photoreceptors in *Drosophila*^1,2^. Further investigation is needed to determine whether if different phototransduction mechanisms are involved in mosquito light preference behavior.

We show the location and oscillation of PER protein at neuronal subgroup level in diurnal and nocturnal mosquitoes. Intriguingly, we find PER protein expression oscillates in anti-phasic manner between diurnal and nocturnal mosquitoes, which could mechanistically underlie diurnal versus nocturnal behaviors. Conceptual support for this finding is that for diurnal versus nocturnal mammals, the circadian clock in non-suprachiasmatic nuclei neurons and periphery tissues cycle in opposite phases between diurnal versus nocturnal rodents and primates^27-29^. It is not known currently what factors contribute to differential phase timing of circadian protein cycling among different tissue and cellular types. Our detailed characterization of light-evoked attraction/avoidance behavior in mosquitoes shows timing features that suggest that these processes are under circadian regulation as we find previously in *Drosophila melanogaster*^1,2^. Experimental verification for this is shown by temporal disruption of light-evoked attraction/avoidance behavior by environmentally shutting down the circadian clock with constant light (LL). A wide range of behaviors in mosquito and other insects are temporally modulated by light, including mating, seeking a blood-meal, biting, oviposition, flight activity, and sleep^9-15^. Light treatments that alter circadian function also disrupts biting, flight activity, and oviposition behaviors^6,7,10,11,14^. By controlling the timing and color of light exposure, we can further specify mosquito species being targeted towards specific manipulation of mosquito behaviors using environmentally friendly light-based approaches.

## Materials and Methods

### Immunocytochemistry

All mosquitoes were reared in standard 12 hr: 12 hr light: dark (LD) schedule at 27°C, and 80% humidity in large cages, with access to 10% sucrose diet. Mosquito brains were dissected 5-10 days post-eclosion. Brains were dissected in 1X PBS, fixed in 4% paraformaldehyde (PFA) for 30 min, washed 3X 10 min in PBS-Triton-X 1%, incubated in blocking buffer (10% Horse Serum-PBS-Triton-X 0.5%) at room temperature before incubation with mouse α-PDF C7, monoclonal (1:10,000) and rabbit α-PER, polyclonal (1:1,000) antibodies overnight in 4° C. Primary antibody incubated brains were then washed 3X 10 min in PBS-Triton-X 0.5% then incubated in goat α-mouse-Alexa- (1:500) and goat α-rabbit-Alexa-594 (1:500) secondary antibodies in blocking buffer overnight in 4°C. Brains were washed 5X 15min in PBS-Triton-X 0.5% before mounting in Vectashield mounting media (Vector Laboratories). Microscopy was performed using Zeiss LSM700 confocal microscope. Fluorescence levels were analyzed using Imaris software (Bitplane). Spherical region of interest was selected for each cell and fluorescence was quantified for each region by the Imaris software. Each species timepont were collected for minimum of three repetitions each. Reported quantification values reflect the average fluorescence intensity levels and error bars indicate S.E.M.

### Light-Induced Attraction/Avoidance Behavioral Assay

All mosquitoes were reared in standard 12 hr: 12 hr light: dark (LD) schedule in 27°C, and 80% humidity in large cages, with access to 10% sucrose diet. Adult mosquitoes (0-5 days post-eclosion) were entrained to LD schedule for minimum of 3 days prior to testing. Individual mosquitoes were each placed into 25mm diameter × 125 mm length pyrex glass tubes (Trikinetics) plugged with “flugs” on either side. Flugs are soaked with 10% sucrose providing a food source, while simultaneously allowing airflow sufficient for multi-day survival of the mosquitoes in the tubes. Tubes containing individual mosquitoes were placed in humidity-, temperature-, and light-controlled incubator and allowed to acclimate for a full day. One half of the tubes were covered with infrared (IR) filters (LEE Filters 4 × 4” Infrared (87C) Polyester Filter), providing the mosquitoes with a choice of a shaded environment (IR filtered) versus light-exposed (not covered with IR filter) during the 12 hrs of light. Philips TL-D Blacklight ultraviolet light (UV) source with narrow peak wavelength of 365 nm and intensity of 400 µW/cm^2^ was used for UV light. Blue (450 nm, Supernight) and red (630 nm, Supernight) LED strips set around 400 µW/cm^2^ was used as blue and red light sources. Additionally, IR LED strips (Infrared 850 nm 3528 LED Strip Light, 78/m, 8mm wide, by the 5m Reel) placed on aluminum heat sink was placed under the entire setup. With each light source, same LD schedule as the LD entrainment schedule prior to experiment was continued to minimize any disturbance to the circadian time. For constant light (LL) light choice assay, the UV light was constantly left on, instead of LD. Webcam (Microsoft Q2F-00013 USB 2.0 LifeCam) took pictures at 5min intervals for 3-5 days of experiment. Each mosquito’s preference in the light-exposed versus shaded side of the tube was analyzed by the ImageJ program. Each experiment was repeated a minimum of three repetition for each group. Preference was averaged for each time point per mosquito and statistical measurements were analyzed by t-test using Microsoft Excel and Sigma Plot.

## Acknowledgments

We thank Annika Barber, Naoki Okamoto, and Anupama Dahanukar for helpful discussions; Guiyun Yan and Adeela Syed for technical support; and Janita Parpana, Eleanor Chan, Duke Park, and Lillian Li for administrative support. TCH is supported by R35 GM127102. LSB acknowledges partial support by an individual NSF pre-doctoral fellowship award. A.R. is founder of Sensorygen Inc, a startup working on natural odorants as insect repellents and food flavorings.

## Author Contributions

L.S.B. conceptualized, designed, acquired, analyzed, interpreted, and drafted this work; C.N. acquired and analyzed data; D.A. analyzed data; T.G. contributed to designing this work; J.A.C. acquired data; A.R. contributed to designing, interpreting, and revising this work; T.C.H. contributed to interpreting and revising this work.

