## Supporting Information for "Circadian regulation of light-evoked attraction/avoidance in day- vs. night-biting mosquitoes"

### Supporting Information Figure Legends

**Supporting Information Figure 1. Schematic of mosquito light-evoked attraction/avoidance behavioral assay.**
**(a)** Schematic of mosquito light-evoked attraction/avoidance preference behavioral assay setup.

**Supporting Information Figure 2. Mosquito light-evoked attraction/avoidance behavior is wavelength-specific.**
**(a-b)** Attraction/avoidance behavior to blue light (450 nm LED, 400 μW/cm$^2$), measured by % preference in blue light-exposed versus shaded environment throughout 12 hr: 12 hr blue light: dark for female *Ae. aegypti* (n=78) **(a)**, female *An. coluzzii* (n=34) **(b)**. **(c-d)** Attraction/avoidance behavior to red light (620 nm LED, 400 μW/cm$^2$), measured by % preference in red light-exposed versus shaded environment throughout 12hr: 12hr red light:dark for female *Ae. aegypti* (n=62) **(c)**, female *An. coluzzii* (n=52) **(d)**. **(e-f)** Average attraction/avoidance behavioral preference to light-exposed versus shaded-environment for daytime **(e)** and nighttime **(f)** in *Ae. aegypti* and *An. coluzzii* female mosquitoes. Data are represented as mean ± S.E.M. *p < 0.05; **p < 0.01; ***p < 0.001 vs. UV.

**Supporting Information Figure 3. Circadian neuronal circuit of diurnal and nocturnal mosquito brains.**
**(a-b)** Representative confocal images of adult female *Ae. aegypti* **(a)** and *An. coluzzii* **(b)** mosquito brains immunocytochemistry stained with α–PER (magenta) and α–PDF (green) antibodies. Similar to *Drosophila*, neurites from PDF$^+$ LNv neurons project dorsally towards the DNs in *Ae. aegypti*. In *An. coluzzii* brains, PDF$^+$ LNv neurites project dorsally towards to the DNs and then extend medially towards the PER$^+$ PI neurons. **(c)** Average number of PERIOD-expressing neurons ±SEM (n= #) in *Ae. aegypti* and *An. coluzzii* female brains, per hemisphere for PDF$^+$ or PDF$^-$ large- and small-LNvs, and per whole brain for DNs, m-ANs, and PI neurons. Female *Ae. aegypti* brains have approximately 8-9 PDF$^+$ l-LNvs and 9-10 PDF$^+$s-LNvs, while female *An. coluzzii* brains have approximately 10 PDF$^+$ l-LNvs and 9-10 PDF$^+$ s-LNvs per hemisphere. Both species of mosquitoes have larger number of LNvs compared to *Drosophila melanogaster*, which has 5-6 l-LNvs and 4-5 PDF$^+$ s-LNvs, but otherwise their neuroanatomical features are highly similar to *Drosophila melanogaster* and other insects. In the lateral ventral region amongst the LNv, there are PER$^+$/PDF$^-$ neurons, again, consistent with a PER$^+$/PDF$^-$ "5$^{th}$ s-LNv" neuron seen in flies. We find approximately 3 PDF$^-$ putative l-LNvs and 6 PDF$^-$ putative s-LNvs in female *Ae. aegypti*, and approximately 3-4 PDF$^-$ putative l-LNvs and 4-6 PDF$^-$ putative s-LNvs in female *An. coluzzii* in each side of the brain.

**Supporting Information Figure 4. PDF$^+$ neurons and in *Aedes aegypti* and *Anopheles coluzzii* female mosquito brains.**
**(a-b)** Representative confocal images of adult female *Ae. aegypti* **(a)** and *An. coluzzii* **(b)** mosquito brains immunocytochemistry stained with α–PDF (green) antibody.

**Supporting Information Figure 5. Oscillatory PER expression of *Aedes aegypti* DNs.**
**(a)** Representative confocal images of adult female *Ae. aegypti* mosquito dorsal neurons (DNs) immunocytochemistry stained with α–PER (magenta) and α–PDF (green) antibodies at ZTs 5,

11, 17, and 23. **(b)** PERIOD expression levels at each ZT for *Ae. aegypti* DNs (ZT 5, n=27; ZT 11, n=17; ZT 17, n=6, ZT 23, n=7). Data are represented as mean ± S.E.M.

**Supporting Information Figure 6. Oscillatory PER expression of *Aedes aegypti* m-ANs.**
**(a)** Representative confocal images of adult female *Ae. aegypti* mosquito mid-anterior neurons (m-ANs) immunocytochemistry stained with α–PER (magenta) and α–PDF (green) antibodies at ZTs 5, 11, 17, and 23. **(b)** PERIOD expression levels at each ZT for *Ae. aegypti* m-ANs (ZT 5, n=27; ZT 11, n=17; ZT 17, n=6, ZT 23, n=7). Data are represented as mean ± S.E.M.

**Supporting Information Figure 7. Oscillatory PER expression of *Anopheles coluzzii* DNs.**
**(a)** Representative confocal images of adult female *An. coluzzii* mosquito dorsal neurons (DNs) immunocytochemistry stained with α–PER (magenta) and α–PDF (green) antibodies at ZTs 5, 11, 17, and 23. **(b)** PERIOD expression levels at each ZT for *An. colluzzii* DNs (ZT 5, n=13; ZT 11, n=31; ZT 17, n=9, ZT 23, n=8)**.** Data are represented as mean ± S.E.M.

**Supporting Information Figure 8. Oscillatory PER expression of *Anopheles coluzzii* PI neurons.**
**(a)** Representative confocal images of adult female *An. coluzzii* mosquito pars intercerebralis (PI) neurons immunocytochemistry stained with α–PER (magenta) and α–PDF (green) antibodies at ZTs 5, 11, 17, and 23. **(b)** PERIOD expression levels at each ZT for *An. colluzzii* DNs (ZT 5, n=13; ZT 11, n=31; ZT 17, n=9, ZT 23, n=8)**.** Data are represented as mean ± S.E.M.

**Supporting Information Movie 1. 3D Rendering of PDF neurons and projections in *Aedes aegypti* female brain.**
3D animation of anti-PDF stained *Aedes aegypti* female brain showing individual z-slice progressing from anterior to posterior, then posterior to anterior, followed by building of z-stack, and Z-stacked brain rotated in multiple directions (+180° around x-axis and back, -180° around x-axis and back, -180° around the y-axis and back, -180° around y-axis and back).

**Supporting Information Movie 2. 3D Rendering of PDF neurons and projections in *Anopheles coluzzii* female brain.**
3D animation of anti-PDF stained *Anopheles coluzzii* female brain showing individual z-slice progressing from anterior to posterior, then posterior to anterior, followed by building of z-stack, and Z-stacked brain rotated in multiple directions (+180° around x-axis and back, -180° around x-axis and back, -180° around the y-axis and back, -180° around y-axis and back).

Supporting Information Figure 1

a

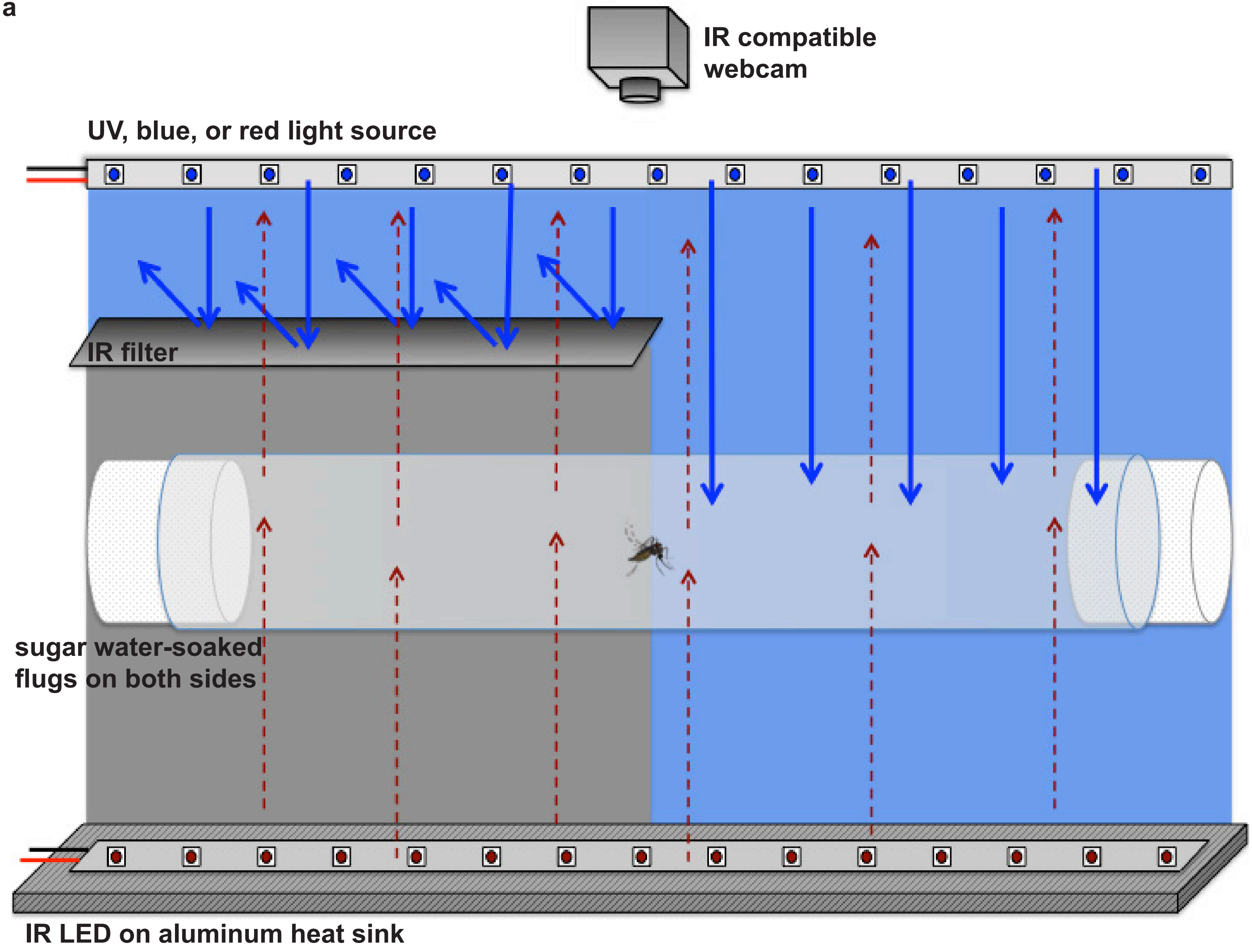

Supporting Information Figure 2

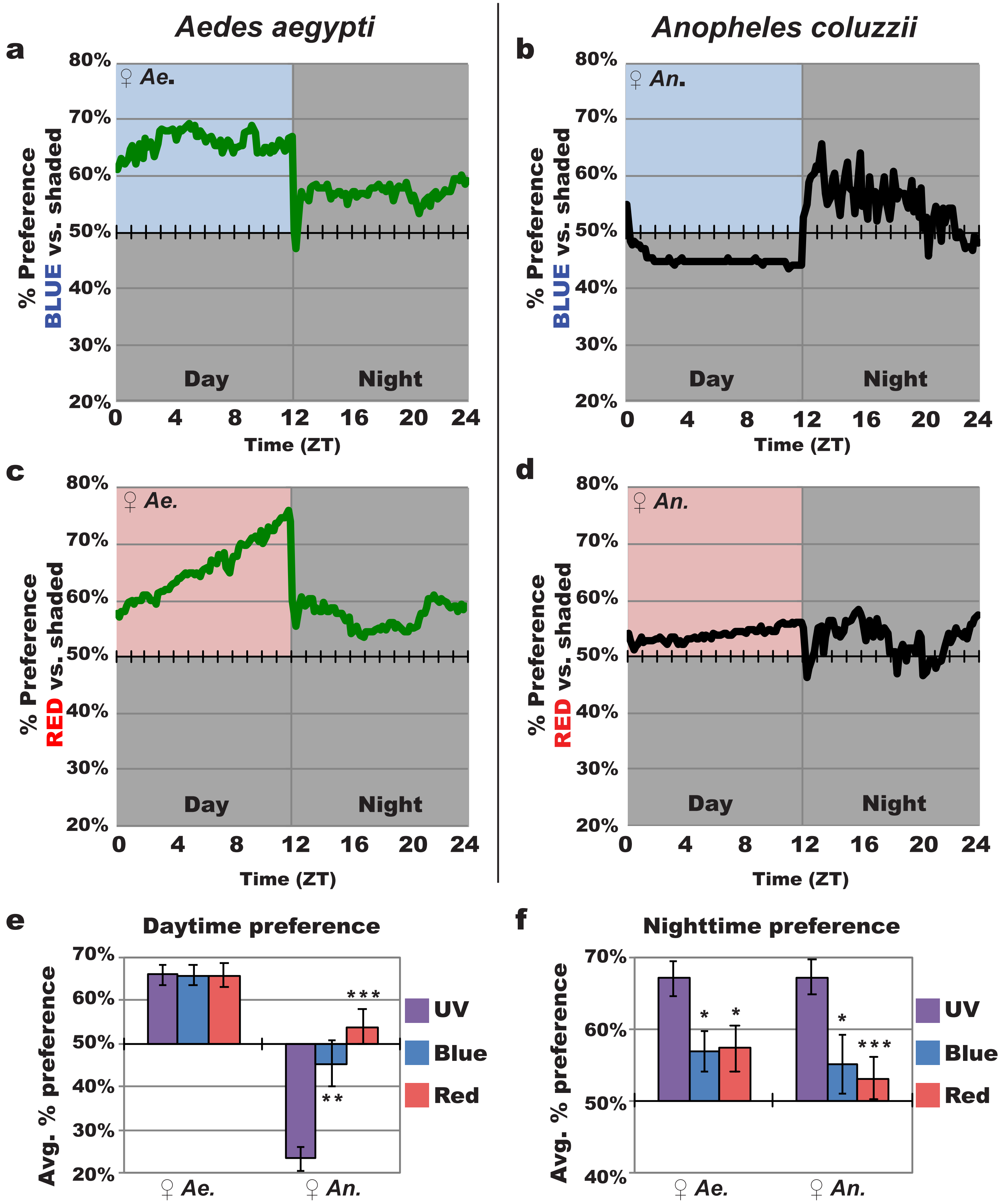

Supporting Information Figure 3

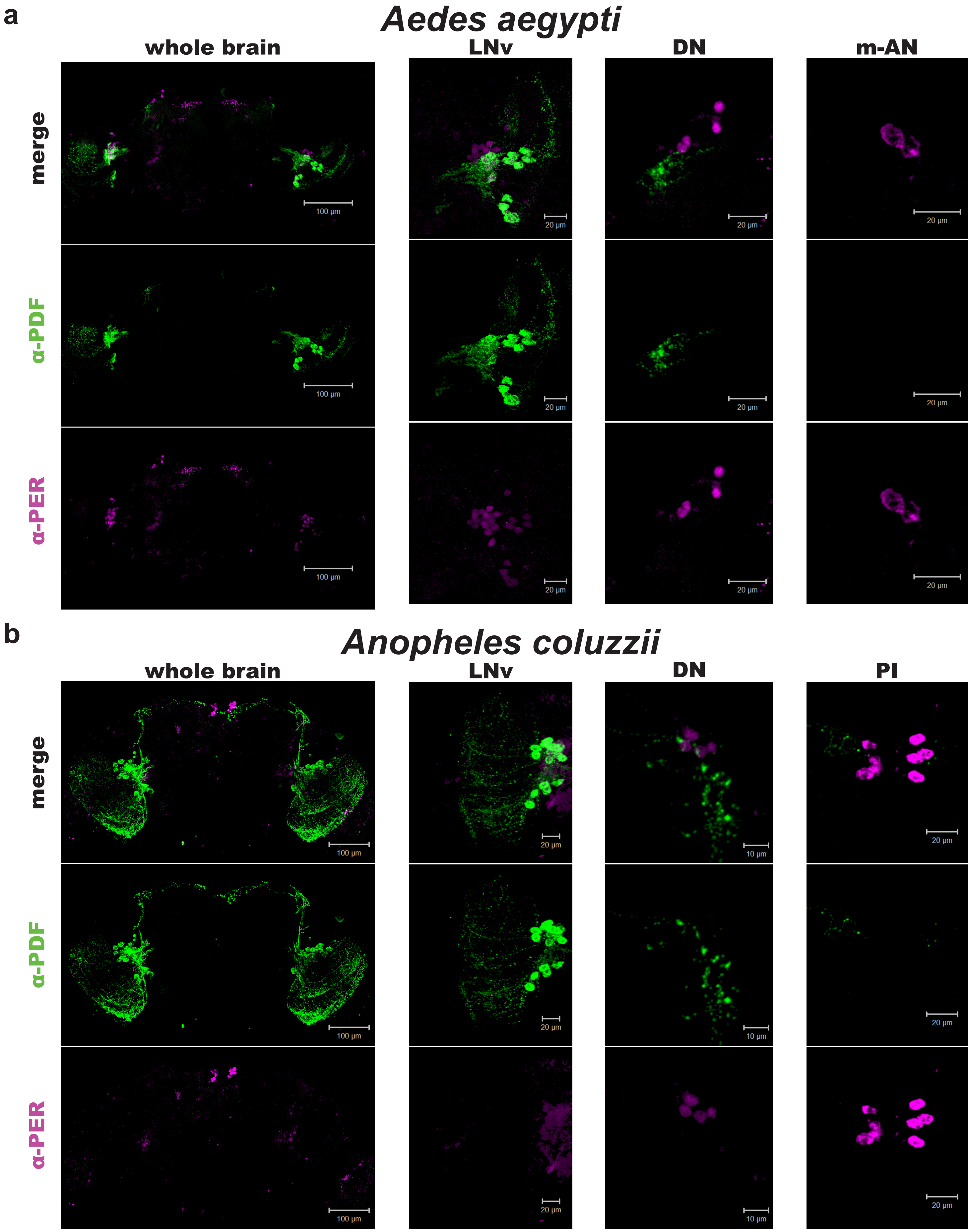


c

| | *Avg. number of neurons per hemisphere* | | | | *Avg. number of neurons per brain* | | |
|---|---|---|---|---|---|---|---|
| | PDF+ l-LNv | PDF+ s-LNv | PDF- l-LNv | PDF- s-LNv | DNs | PI Neurons | m-ANs |
| *Aedes aegypti* | 8.6 ± 0.4 (n= 18) | 9.3 ± 0.5 (n= 18) | 3.1 ± 0.4 (n= 19) | 5.8 ± 1.1 (n= 19) | 4.1 ± 0.3 (n= 30) | - | 4.8 ± 0.4 (n= 31) |
| *Anopheles coluzzii* | 10 ± 0.5 (n= 20) | 9.8 ± 0.5 (n= 21) | 3.5 ± 0.7 (n= 8) | 5.2 ± 0.8 (n= 8) | 3.3 ± 0.4 (n= 26) | 7.3 ± 0.1 (n= 22) | - |

Supporting Information Figure 4

**a**

***Aedes aegypti***

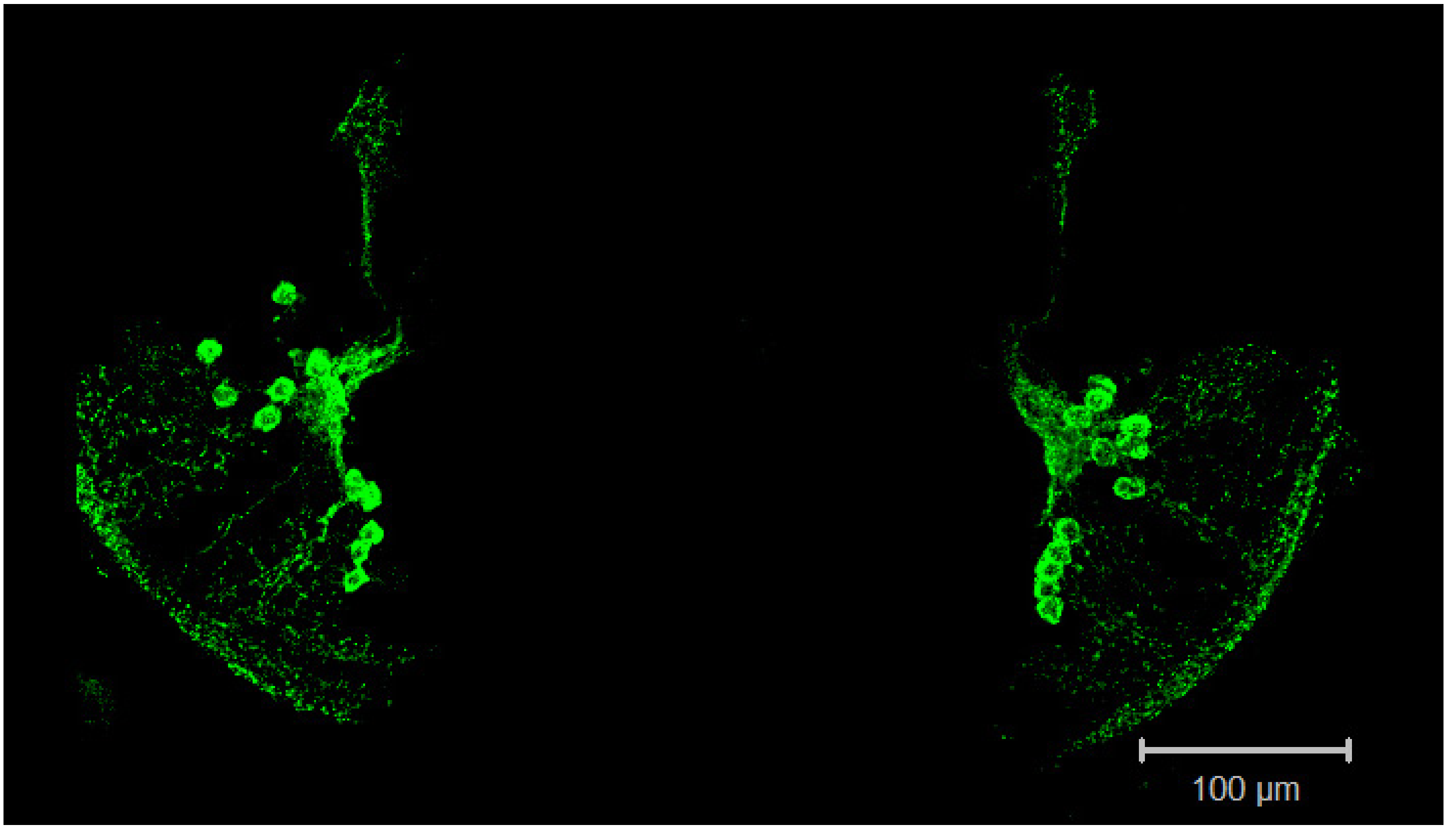


**b**

***Anopheles coluzzii***

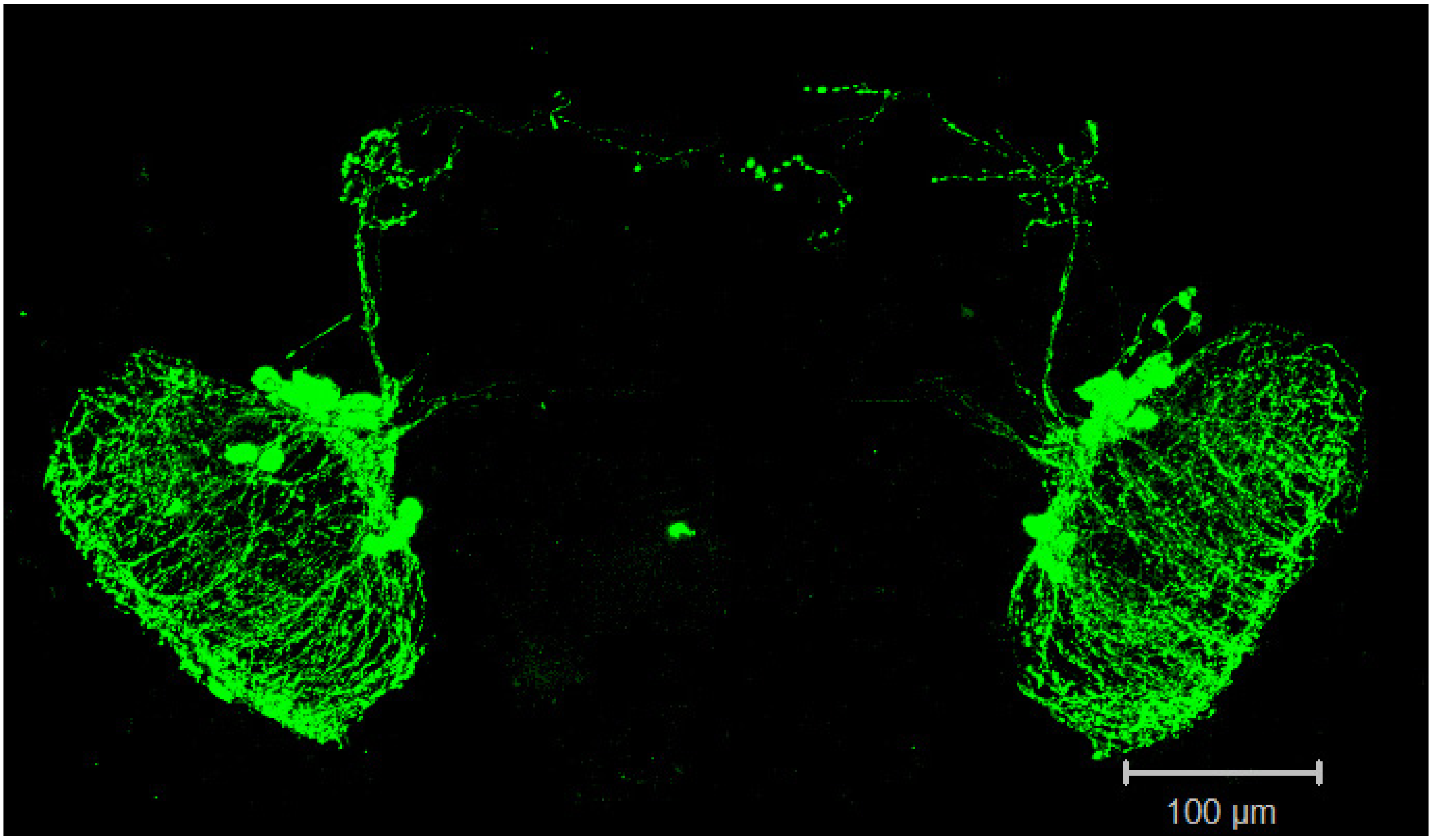

Supporting Information Figure 5

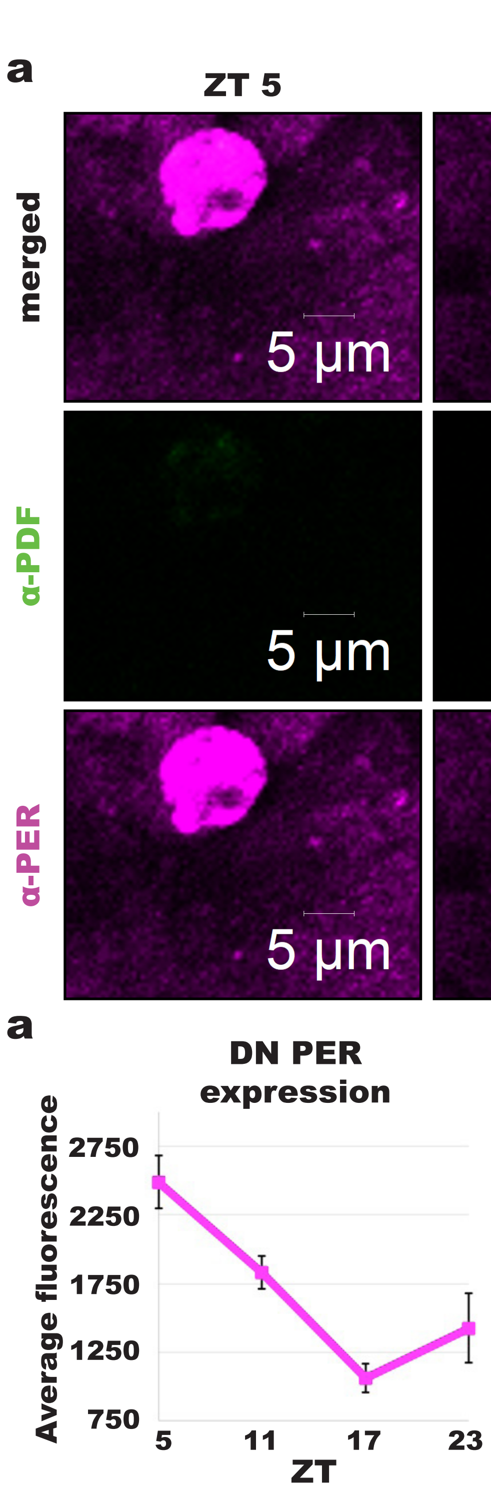

Supporting Information Figure 6

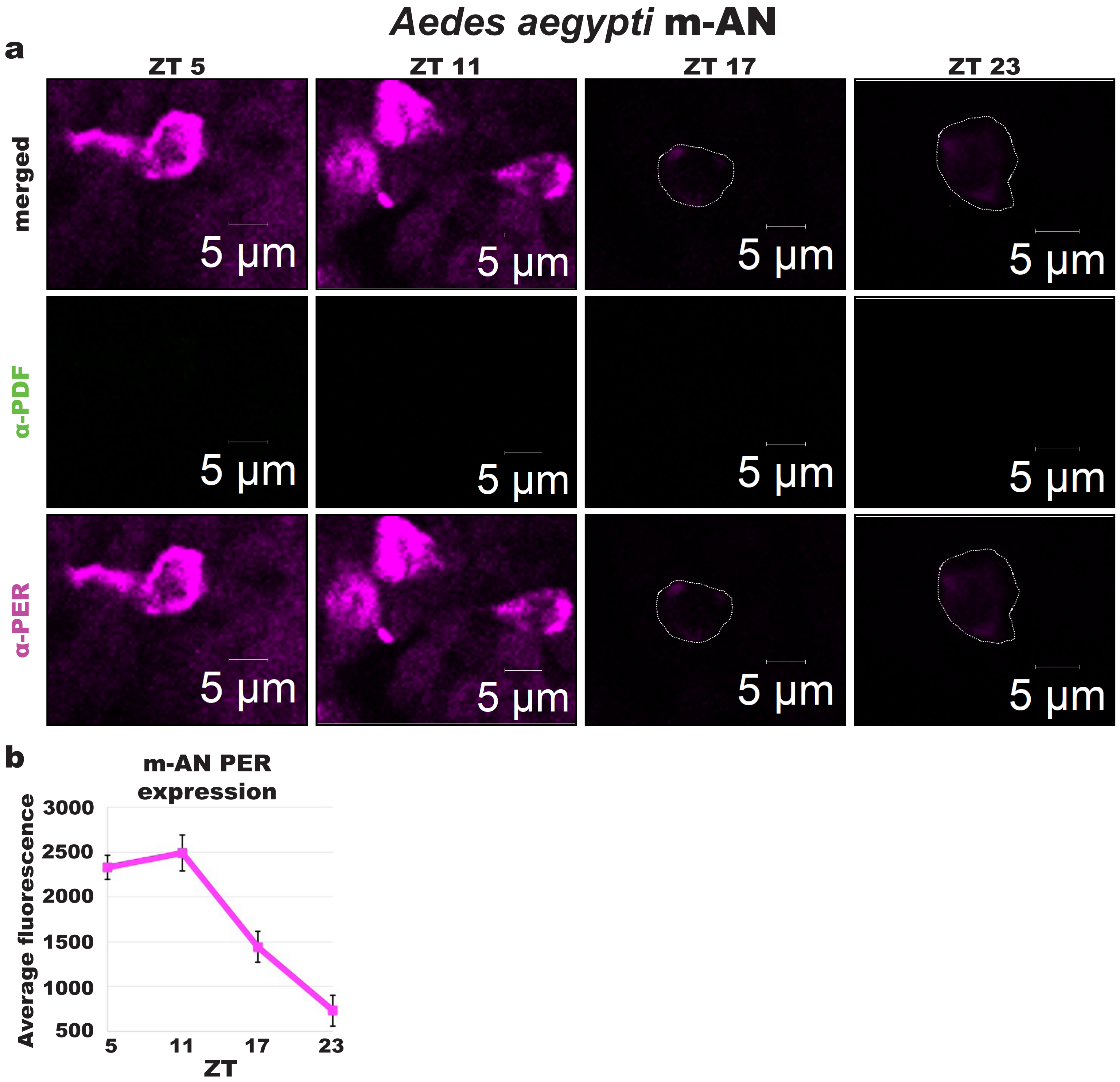

Supporting Information Figure 7

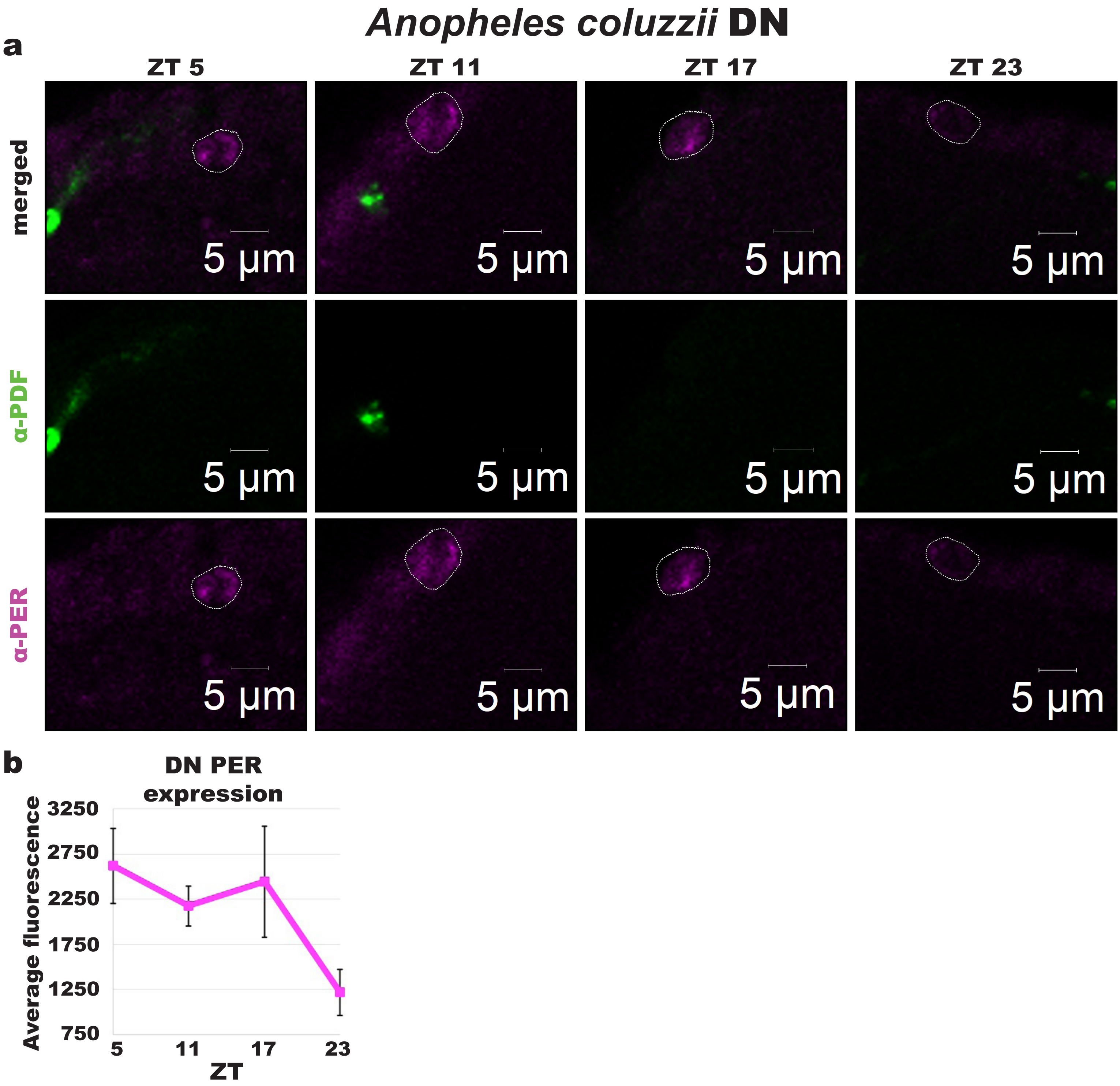

Supporting Information Figure 8

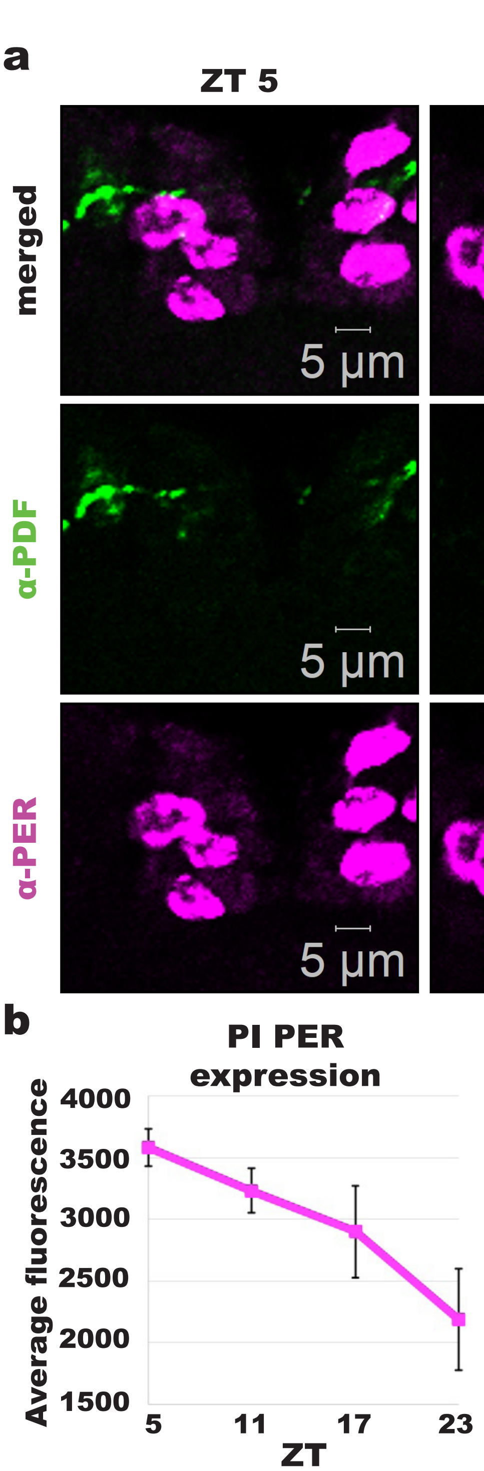

Supporting Information Movie 1

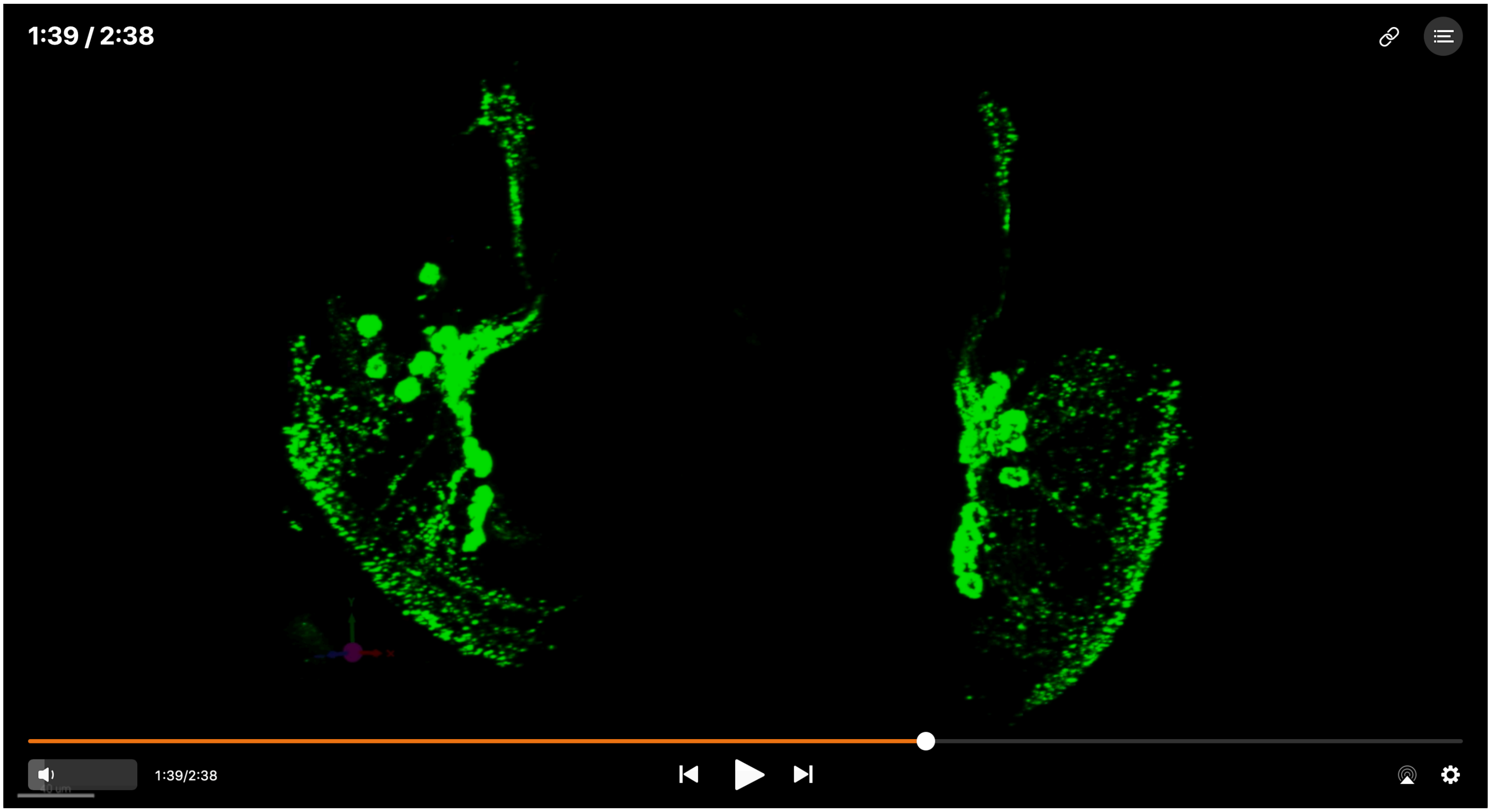

Supporting Information Movie 2

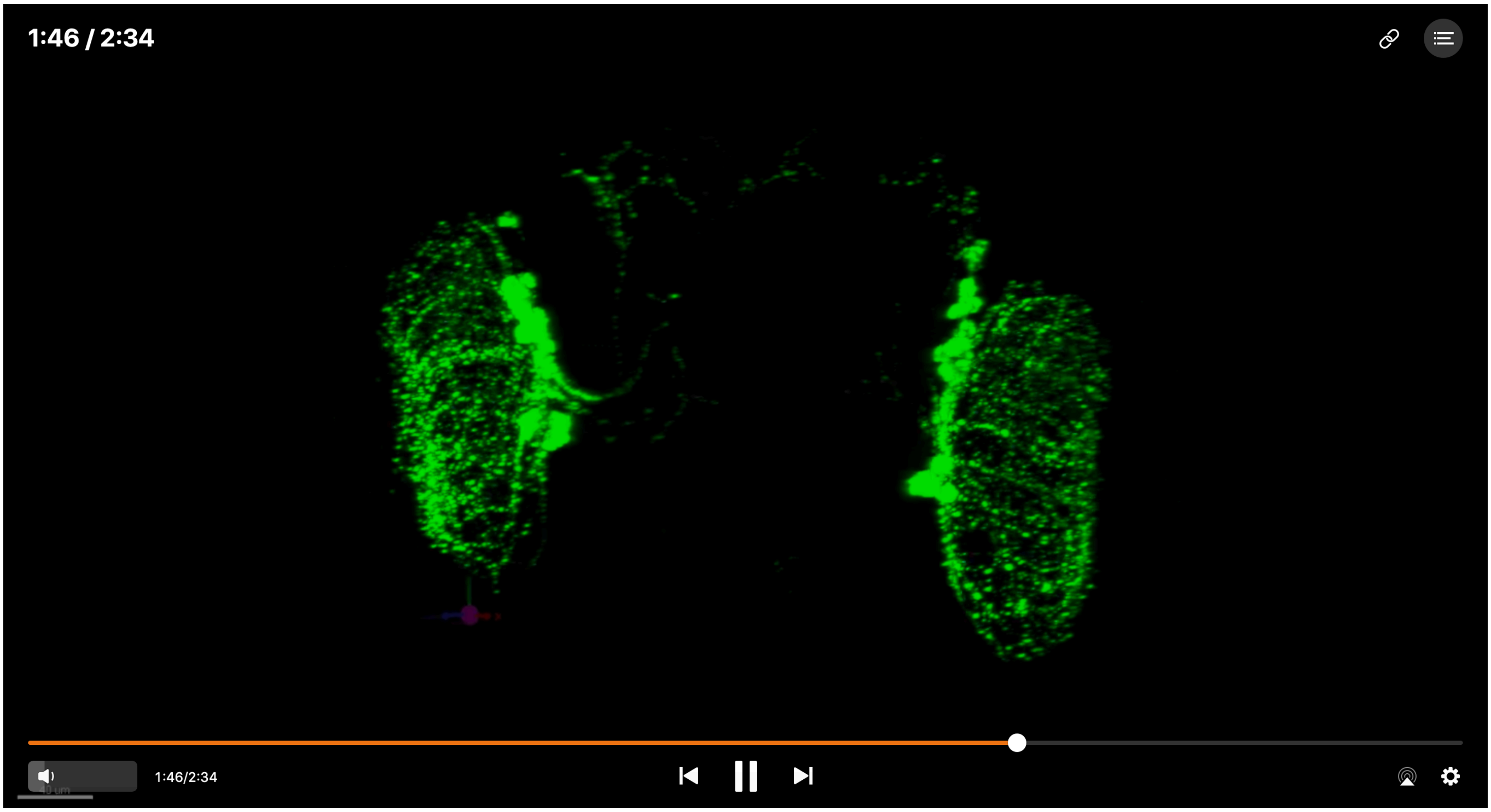
